# Fast and Accurate LSTM Meta-modeling of TNF-induced Tumor Resistance In Vitro

**DOI:** 10.1101/2024.08.12.607535

**Authors:** Marco P. Abrate, Riccardo Smeriglio, Roberta Bardini, Alessandro Savino, Stefano Di Carlo

## Abstract

Multi-level, hybrid models and simulations, among other methods, are essential to enable predictions and hypothesis generation in systems biology research. However, the computational complexity of these models poses a bottleneck, limiting the applicability of methodologies relying on large number of simulations, such as the Optimization via Simulation (OvS) of complex biological processes. Meta-models based on approximate surrogate models simplify multi-level simulations, maintaining accuracy while reducing computational costs. Among Artificial Neural Networks (ANNs), Long Short-Term Memory (LSTM) networks are well suited to handle sequential data, which often characterizes biological simulations. This paper presents an LSTM-based surrogate modeling approach for multi-level simulations of complex biological processes. Validation relies on the simulation of Tumor Necrosis Factor (TNF) administration to a 3T3 mouse fibroblasts tumor spheroid based on PhysiBoSS 2.0, a hybrid agent-based multi-level modeling framework. Results show that the proposed LSTM meta-model is accurate and fast compared with the simulator. In fact, it infers simulated behavior with an average relative error of 7.5%. Moreover, it is at least five orders of magnitude faster. Even considering the cost of training, this approach provides a faster, more accurate, and reusable surrogate of multi-scale simulations in computationally complex tasks, such as model-based OvS of biological processes.

## I. Introduction

Multi-level, hybrid models and simulations are essential to support research in systems biology, allowing to predict behaviors of biological systems and generate hypotheses for further validation [1], [2]. In addition, they can support Design Space Exploration (DSE) of complex biological processes, that is, the process of identifying optimal design solutions that meet specific requirements by algorithmically evaluating a range of potential design options [3]. Optimization via Simulation (OvS), the iterative process of determining the best configuration of a simulated system in order to optimize one or more objective functions, supports the DSE of complex biological processes [4] such as biofabrication protocols in tissue engineering and regenerative medicine [5]–[7], and drug administration schemes in pharmacology [8]. This task is based on a continuous exchange between the optimization task, which selects potential candidates for the optimal solution, and the simulation, which evaluates these candidates [9].

In particular, optimizing drug treatments and efficiently screening drug effects are essential for improving clinical outcomes and extending patients’ life expectancy. However, the complex interplay of biological processes and environmental factors underlying the emergence of tumor resistance is difficult to factor in treatment optimization [8]. Multi-level hybrid models and simulations are valuable tools for studying these phenomena, as they integrate molecular, cellular, and intercellular processes, providing insights into the dynamics of cancer drug resistance and the development of optimized treatment regimens [8]. Yet, they require high computing power, posing a bottleneck in the execution of OvS algorithms, which typically require a large number of simulations for DSE.

To tackle this limitation, the use of meta-models in this context has the potential to accelerate DSE by reducing computational costs and time linked to simulation. Meta-modeling approaches, based on the development of surrogate models, have the potential to approximate multi-level simulations while maintaining accuracy and reducing computational costs [10]. Common meta-models base on polynomials, kriging, and Artificial Neural Networks (ANNs) [4], [11], [12]. Among ANN models, Recurrent Neural Networks (RNNs) and Long Short-Term Memory (LSTM) networks are designed for handling time series [13]; they are, therefore, good candidates to support the meta-modeling of simulated processes. This paper proposes an LSTM-based surrogate model for multilevel simulations of complex biological processes, yielding accurate inference of the simulation evolution throughout its duration. The validation of such a model relies on a *in vitro* pharmacology use case, where the objective is to design drug administration schemes to optimize pharmacological effects and later translate them clinically. In particular, the use case is a Tumor Necrosis Factor (TNF) stimulation of 3T3 mouse fibroblasts tumor spheroid case study [14], [15]. This reference case study is simulated using PhysiBoSS 2.0 [16], a hybrid agent-based modeling framework. PhysiBoSS simulates signaling and regulatory networks within individual cell agents, expanding PhysiCell [17] by enabling intracellular simulations with MaBoSS [18] to study the interplay between the microenvironment, signaling pathways, and cellular dynamics.

This work lays the groundwork for developing a tool that biologists can use for fast and accurate prediction of systems behavior and efficient DSE of complex biological processes. Results show the LSTM meta-model accurately reproduces the behavior of the simulated use case, reducing the computational time required for prediction by at least five orders of magnitude compared to simulation while preserving high accuracy in predicting the evolution of the system behavior. This shows the potential of the LSTM approach for time-series-based metamodeling of complex simulated biological behaviors, paving the way for efficient biological DSE.

The paper is organized as follows: Section II presents the general context of surrogate modeling for DSE of complex processes, and the technological approaches based on RNNs in particular; Section III introduces the proposed methodology; Section IV shows the results obtained for the meta-modeling of the PhysiBoSS 2.0 TNF tumor spheroid use case; Section V analyzes potentials and limitations of the proposed approach, as well as future research directions.

## II. Background

Surrogate models, also known as meta-models, are crucial to reduce the computational costs of OvS [10], [12], allowing efficient exploration and optimization of a biological process design [19]. They can be classified based on their level of interpretability, the science of comprehending the actions of a model [20], and the amount of system knowledge they incorporate [21]. White-Box models are fully transparent and interpretable. Black-Box models are highly flexible and powerful, they operate with minimal understanding of the system’s internal processes, focusing on input-output data. Black-Box models are capable of modeling complex, non-linear relationships but need dedicated effort to increase their transparency and interpretability. Gray-Box models combine elements of White-Box and Black-Box models, offering partial interpretability while leveraging data-driven methods to manage complexity and uncertainty.

Among black-box models, RNNs are well-suited as surrogate simulation models because they can handle time series to represent a process [22]. RNNs are designed to handle time series by retaining information from previous inputs in their internal state. This capability makes them suitable for modeling processes where the order of events is crucial. However, traditional RNNs can struggle with learning longterm dependencies due to issues like vanishing gradients [23]. LSTMs [24] overcome the limitations of traditional RNNs by incorporating gating mechanisms that regulate the flow of information [23]. The key insight in the LSTM design was to incorporate non-linear, data-dependent controls into the RNN cell, which can be trained to ensure that the gradient of the objective function with respect to the state signal (the quantity directly proportional to the parameter updates computed during training by gradient descent) does not vanish [25]. This design enables LSTMs to learn long-term dependencies effectively, making them a perfect candidate for modeling complex sequential processes in biological systems, where understanding the evolution of states over time is essential [26]. Despite belonging to the Black-Box model category, LSTMs are interpretable through visualization-based approaches like attention mechanisms and saliency maps by highlighting influential parts of the input sequence. LSTM surrogate models have been successfully employed in computational biology to replicate the behavior of a Stochastic Differential Equation (SDE) model describing the MYC/E2F transduction pathways in cell-cycle progression [27], to Partial Differential Equations (PDEs) simulation of complex spatio-temporal problems [28], to Agent-based models (ABMs) of cytotoxic T-lymphocyte and cancer cells interactions [29] as well as for oxygen uptake prediction [30]. They have also been used to solve more general classification and regression problems, such as PM2.5 prediction [31] and traffic flow prediction [32].

The two major limitations of ANN-based surrogate models are the computational cost of training, and generalizability across systems as most approaches are model-specific. To take a first step towards improved generalizability, this work introduces a surrogate model of PhysiBoSS 2.0 [16] - a robust, multi-level, and hybrid simulation framework supporting the simulation of a broad range of biological systems. Seamless integration with this general tool sets the stage for developing new metamodeling processes for systems biology simulations that can handle a wide range of use cases [33].

## III. Methods

The proposed meta-model is presented through a running example aiming of reproducing the behavior of cells under TNF stimulation in a 3T3 mouse fibroblasts tumor spheroid, simulated with PhysiBoSS 2.0. In the selected use case, the tumor cells undergo periodic TNF stimulations that possibly trigger them to transition from a proliferative state towards necrotic or apoptotic states [8]. The trends of the number of cells in these states can change because of three different input parameters: the *Pulse period*, the *Pulse duration*, and the *TNF concentration*. In our use case, the simulator replicated the system’s behavior over a full day (1440 minutes) in an invitro system, taking around 7 seconds of Central Processing Unit (CPU) time. The meta-model predicts the number of alive (*N*_alive_), apoptotic (*N*_apoptotic_), and necrotic (*N*_necrotic_) cells every 60 minutes throughout the entire simulation.

### A. LSTM meta-model

To handle sequential and temporal data and to avoid problems related to vanishing gradients, the LSTM architecture was implemented [23].

The architecture we developed (Figure 1) is implemented with Pytorch [34], and comprises three main components:

**Fig 1:**
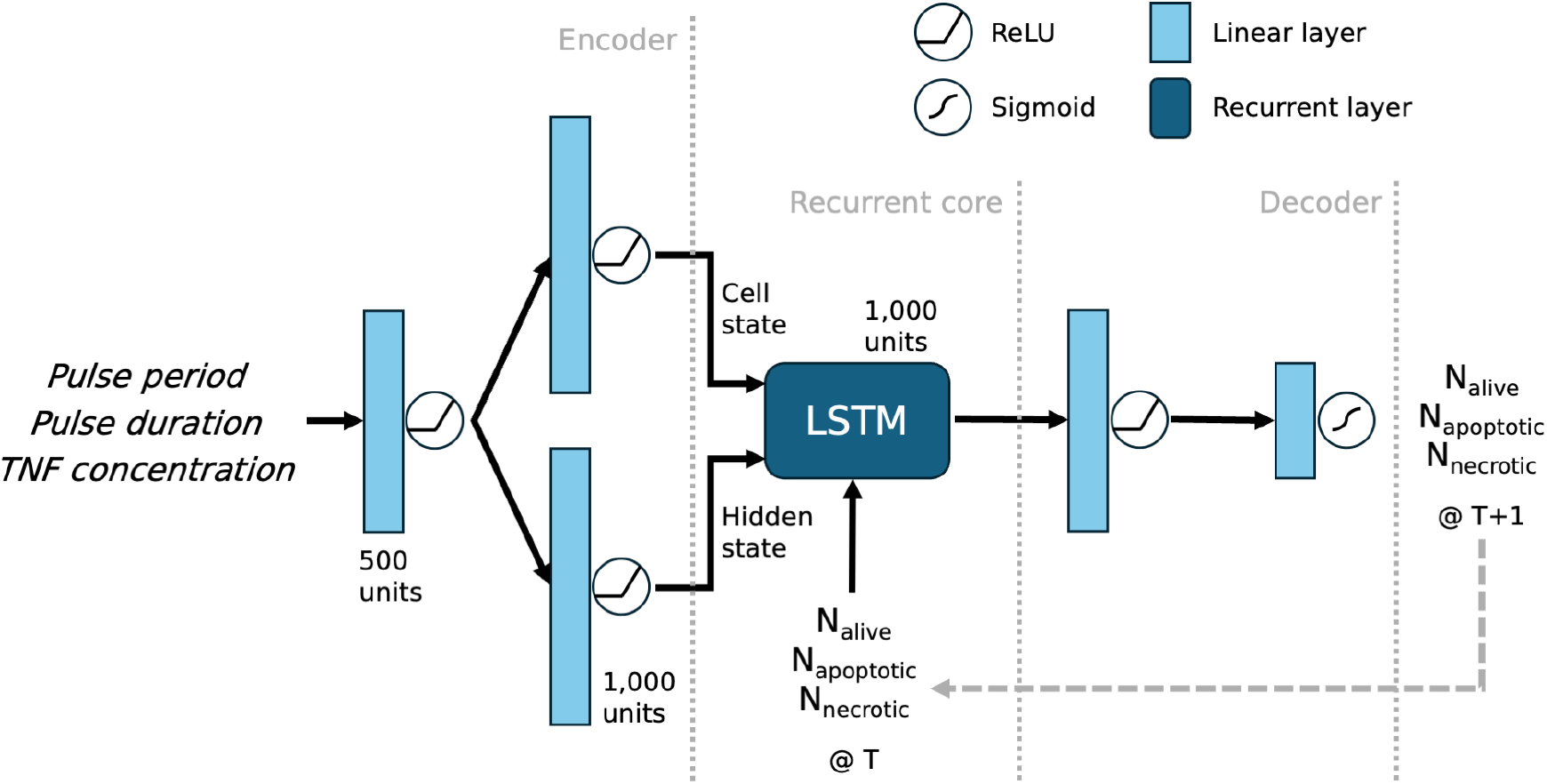
Schematic of the LSTM meta-model architecture. It is composed of an encoder that transforms the input parameters into a latent representation suitable for the recurrent network, a recurrent core that is a single layer LSTM network, and a decoder that processes the output of the recurrent core into the predicted number of cells. The dashed arrow represents inference.

1. The *Encoder* (Figure 1, left) transforms the input parameters (*Pulse period, Pulse duration*, and *TNF concentration*, see Section III-B), which are previously normalized through min-max scaling, into a latent representation suitable for the recurrent network. It consists of a series of linear layers with Rectified Linear Unit (ReLU) activation functions. Two final encoder layers are in charge of splitting the output, one feeding into the LSTM hidden state and the other into the LSTM cell state.
2. The *Recurrent core* (Figure 1, middle) is a single layer LSTM network with dropout regularization responsible for processing the sequential data by taking the encoded parameters and the number of cells from the previous timestep as inputs.
3. The *Decoder* (Figure 1, right), with a similar architecture to the encoder, processes the output of the recurrent core into the predicted number of cells for the following timestep. The final layer uses a Sigmoid activation function to ensure the output is constrained between 0 and 1. This value is then converted to the number of cells by multiplying it by the normalization factor.

### B. Dataset Construction

The training of the meta-model is based on a subset of all possible combinations of input parameters to guarantee sufficient coverage of all relevant cases. Among several input parameters, we chose to explore the values of three salient parameters to extract a dataset that best represents the wide design space of the solutions. We picked biologically plausible ranges [8] to build their value combinations:

1) *Pulse period*: the period of TNF pulses administration along the simulation, ranging between 5 and 800 minutes;
2) *Pulse duration*: the duration of each pulse of TNF administration, ranging between 5 and 200 minutes;
3) *TNF concentration*: the concentration of the TNF stimulus at each pulse, ranging between 0.1009 and 1 TNF*/µm*^3^;

We extracted 1,822 combinations of *Pulse period, Pulse duration*, and *TNF concentration* by considering all possible combinations of 11 equally spaced values for each parameter ranges, along with additional combinations involving the 10 middle values. This approach increases the dataset size and improves training performance. The starting *tumor radius* of the spheroid, which directly defines the initial number of cells, was set to 50, 100, 275, or 400 *µm*. In fact, with a radius of less than 50 *µm*, the number of initial cells is too small to produce a significant experiment, while a radius of more than 400 *µm* requires an impractically long simulation time. For each *tumor radius* value, multiple simulations were executed using the selected combinations of input parameters. The simulations were grouped according to the initial *tumor radius* because the number of cells increases significantly with it, causing normalization issues in the training set. Hence, a separate LSTM meta-model was trained on each group, as described in Section IV-A.

To summarize, this process resulted in the creation of four datasets (one for each *tumor radius*), with each containing 1,822 simulations. *N*_alive_, *N*_apoptotic_, and *N*_necrotic_ were recorded every 60 simulated minutes, resulting in 24 timesteps per simulation covering all 1,440 simulated minutes.

### C. Pre-processing

Each of the four datasets was split into 1,603 simulations (around 88% of the available data) for the training set and 219 (around 12%) for the validation dataset. The status of the cell types was pre-processed to better train the network. Specifically, *N*_alive_, *N*_apoptotic_, and *N*_necrotic_ were normalized separately for each dataset using min-max scaling:

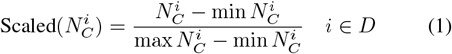

where 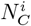 represents the number of cells in state *C* of the *i*^*th*^ dataset, and *D* represents the datasets. Finally, the training simulations were grouped into 7 batches of 229 simulations each, while the validation dataset consisted of 3 batches of 73 simulations each.

### D. Metrics

#### a) Training and validation metrics

The meta-models were trained to minimize the average error of a batch corresponding to the sum of the Mean Squared Errors (MSEs) between the numbers of cells predicted by the LSTM and the true ones over all cell types prior to re-normalization of the values. More in detail, given the number of cells per type *N*_*C*_ and a prediction 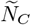, the objective function *L* is calculated as:

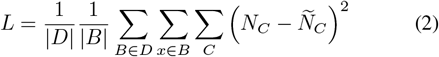

where *D* indicates the datasets, *B* the batches, and *C* the cell type - alive, apoptotic, or necrotic. This MSE is also calculated on the validation dataset to evaluate the model’s performance.

#### b) Evaluation metrics

Since this work is mainly focused on the reproduction of the entire evolution of the simulations, we explored the performance trend over the entire simulated time. Thus, we computed the relative error *R* of the predictions on the total number of cells at simulated timestep *T* as:

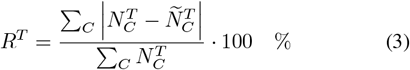

where *C* is the cell type. To evaluate the accuracy of the prediction for each cell type *C*, we also evaluated the respective Mean Absolute Error (MAE) of the prediction at each simulated time *T* as:

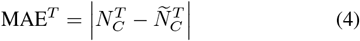

Finally, we selected a random simulation for each batch to qualitatively compare the trend predicted by the meta-model against the actual trend of the simulation.

## IV. Results & Discussion

The CPU used for the simulations is a 12th Gen Intel(R) Core(TM) i9-12900K with 24 threads and a maximum frequency of 5.2 GHz, while the Graphical Processing Unit (GPU) used for the training of the model is an NVIDIA RTX A4000 with 16 GB of memory. Results show that the proposed LSTM meta-modeling approach is fast and accurate in reproducing simulation results under a broad range of TNF administration schemes. Indeed, the trained meta-model successfully replicates the behavior of all simulations outside of the training set. It is accurate enough to be considered a faithful reproduction of the PhysiBoSS 2.0 simulation under different TNF injection patterns and for multiple tumor radii. Moreover, meta-model-based inference is five orders of magnitude faster than simulation.

### A. Training

LSTM training relies on Back Propagation Through Time (BPTT) and uses full simulation sequences, represented with a window of 24 timesteps (see Section III-B). As reported in Section III-B, we generated four specialized models, each tailored to accurately replicate the behavior associated with its specific *tumor radius* condition. For all models, the objective function is the sum of the Mean Squared Error (MSE) between the number of cells predicted by the model and the true ones (see III-D, Equation 2). Trainings lasted 3,000 epochs. During each epoch, the LSTM iteratively processed batches of training data, updating its parameters to minimize the discrepancy between predicted and actual *N*_alive_, *N*_apoptotic_ and *N*_necrotic_ leveraging on the *RMSProp* optimizer to adjust the model’s weights according to the computed gradients. The learning rate was set to 10^*-*6^ and reduced by a factor of ten if the validation loss plateaus for more than 25 epochs, to facilitate convergence. Figure 2 depicts the trends of the training and validation losses. Throughout the training, the loss decreases reaching a minimum without over-fitting the data. This is likely because the implemented model is not overly complex, given the low number of linear layers, the small dimension of the LSTM hidden layer, and the dropout regularization.

**Fig 2:**
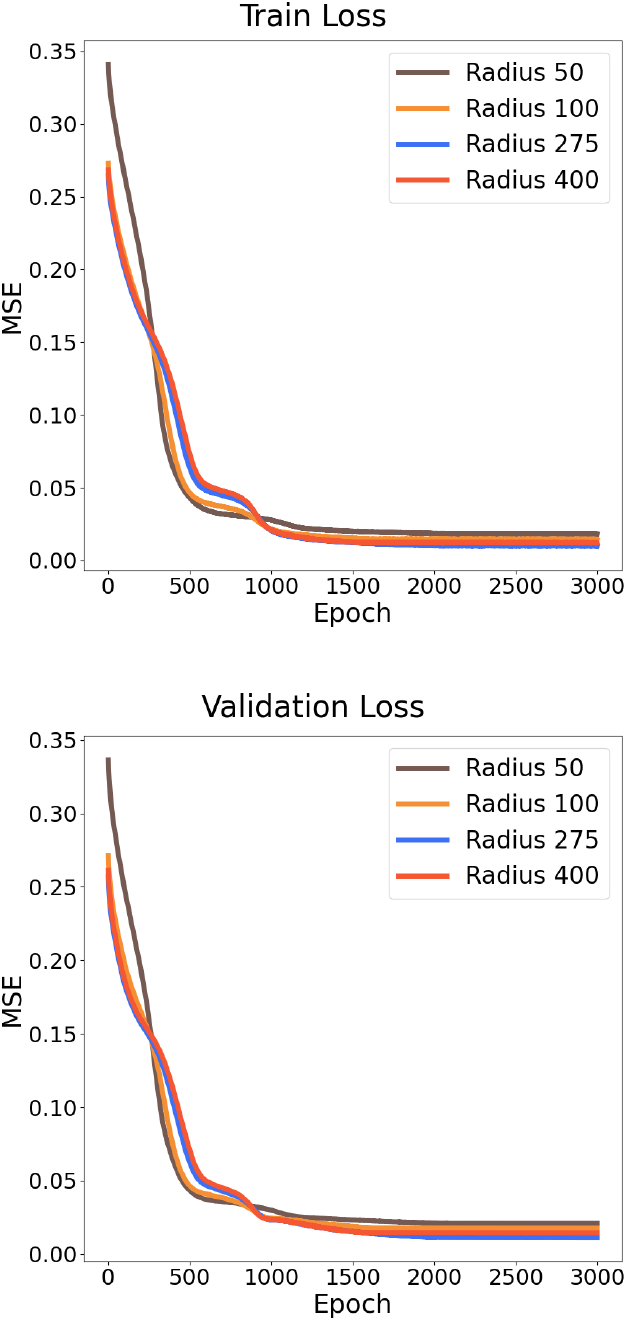
Mean Squared Errors (MSEs) train (top panel) and validation loss (bottom panel) trends throughout the metamodel’s training for the four *tumor radius* value conditions.

### B. Prediction performance

Besides evaluating training performance, this work explores validation metrics (see Section III-D) to evaluate the accuracy of the trained models throughout the entire simulation duration. Figure 3 shows the relative MAE (see III-D, Equation 3) on the number of cells predicted by the model over the entire simulation period across the different *tumor radii*. The results indicate that trends of relative MAE varies significantly with the *tumor radius*. As highlighted in the later portion of the simulated time, maximum relative MAE is higher for smaller radii and lower for larger ones. For all four conditions, the relative MAE increases with the simulation time over the validation dataset. In fact, working with a recurrent model, the error relative to each predicted step propagates to the subsequent steps, causing error accumulation along inference steps. In more detail, the model exhibits a rapid increase in the relative error for radius 50 just before halftime of the simulations, peaking at approximately 25%. A similar trend is observed for radius 100, but the error stabilizes around 15% towards the end of the simulation period. In contrast - for the two larger tumor radii - the LSTMs maintains a lower and stable relative error, peaking at around 5%.

**Fig 3:**
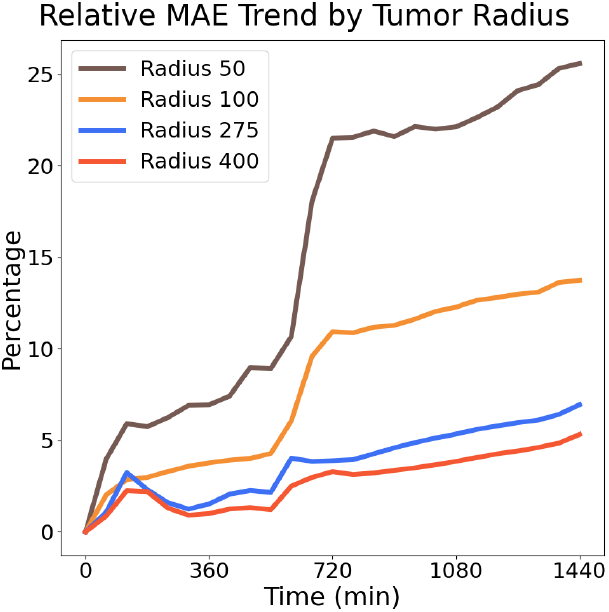
Trends of the relative MAE on the number of cells predicted by the meta-model during simulation time. Each line represents the error for each value of the *tumor radius*.

Furthermore, we analysed the MAE trends for *N*_alive_, *N*_apoptotic_, and *N*_necrotic_ over the simulation period for each *tumor radius* (see Figure 4). The main trend is a reduction of the errors by increasing the *tumor radius*, as previously observed for the relative error. The results highlight distinct behaviors in the prediction errors of each cell type. For apoptotic cells - when the *tumor radius* is 50 or 100 *µm* - the MAE is high during the first half of the simulation period, indicating apoptosis could be accurately predicted once the initial transient phase is over (see Figure 4, top). When using higher *tumor radii, N*_apoptotic_ is predicted with small errors throughout the entire simulation. For *N*_alive_ and *N*_necrotic_, the MAE trend increases with simulation time. This highlights that capturing the behavior of larger *tumor radii* is easier, may be due to more pronounced patterns in the data for larger *radii*, which the model can learn more effectively. This could be linked to the prevalence of stochastic effects over the smaller numbers of cells corresponding to smaller *radii*.

**Fig 4:**
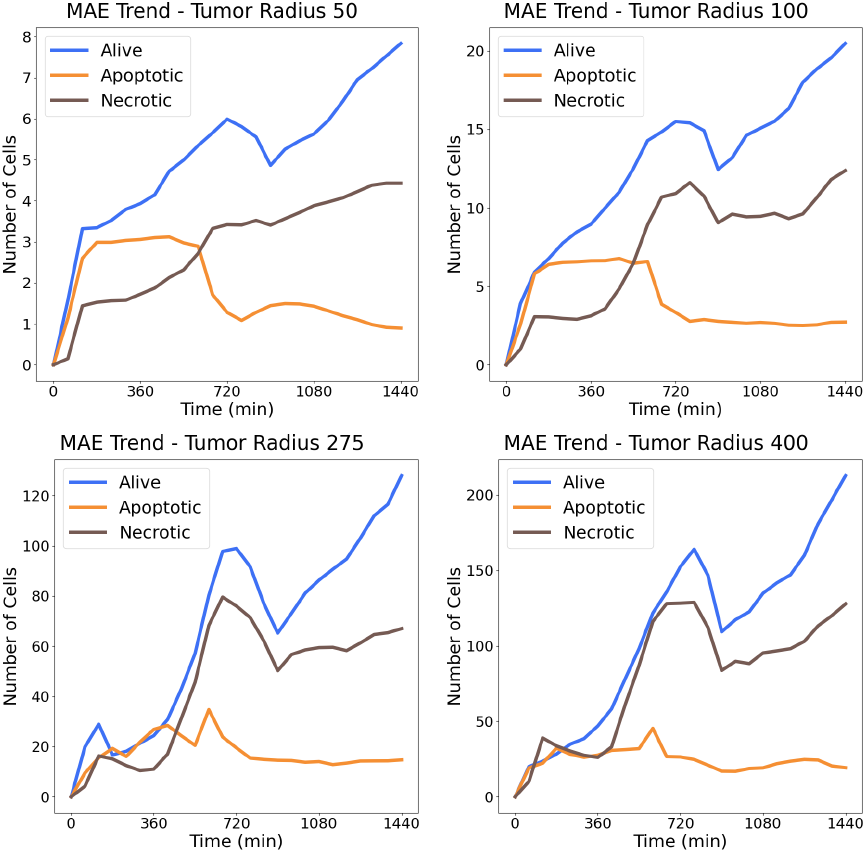
Trends of the MAE on the number of cells predicted by the meta-models during simulation time for all cell types. Plots show the errors for different values of the *tumor radius*: *(top left)* 50, *(top right)* 100, *(bottom left)* 275, and *(bottom right)* 400. Note the change in scale of the *y*-axis to highlight the trends.

Finally, Figure 5 shows four examples of trends of *N*_alive_, *N*_apoptotic_, and *N*_necrotic_ over the simulation period predicted by the meta-model during simulation time. Continuous line represents ground truths (PhysiBoSS 2.0 simulations), while dashed lines represent predictions. Plots show examples for different values of the *tumor radius*: *(top left)* 50, *(top right)* 100, *(bottom left)* 275, and *(bottom right)* 400. The metamodel predictions are a very accurate representation of the cell states’ dynamics produced by the simulator.

**Fig 5:**
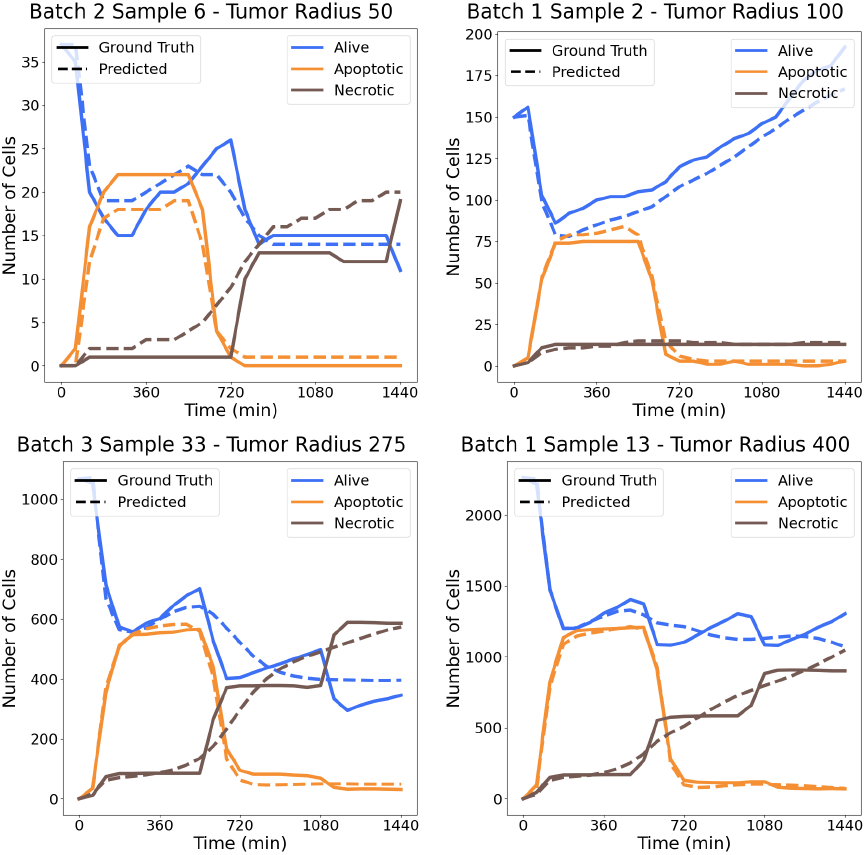
Examples of trends of *N*_alive_, *N*_apoptotic_, and *N*_necrotic_ over the simulation period predicted by the meta-model during simulation time. Continuous lines represent ground truths (PhysiBoSS 2.0 simulations), while dashed lines represent predictions. Plots show examples for different values of the *tumor radius*: *(top left)* 50, *(top right)* 100, *(bottom left)* 275, and *(bottom right)* 400.

**TABLE I:** Numerical comparison between simulations and meta-model’s inference times for the different values of *tumor radius*.

| Tumor radius | Simulation time | Meta-model time |
| --- | --- | --- |
| 50 $\mu m$ | 2.55 s | 0.04 ms |
| 100 $\mu m$ | 3.02 s | 0.04 ms |
| 275 $\mu m$ | 7.38 s | 0.04 ms |
| 400 $\mu m$ | 13.45 s | 0.04 ms |

### C. Computational time reduction

Once trained, the meta-model offers significant time benefits and reduces the usage of computational resources when running simulations. This efficiency comes at the cost of only a negligible drop in accuracy, making it a highly effective tool for large-scale or repeated simulations (see Section IV-B). Specifically, once training set generation is complete, the training of the neural network takes around 3 minutes and 30 seconds of GPU time, independently of the *tumor radius*. The inference has constant time cost, being almost instantaneous (around 8 milliseconds for the entire validation dataset, composed of 219 simulations). Conversely, the time required to execute a simulation with PhysiBoSS 2.0 changes as a function of the *tumor radius*, since higher numbers of simulated cells require larger computational resources. Table I shows the CPU times compared to the meta-model’s inference time.

This significant time improvement is evident, with our metamodel inferring simulation evolution at least five orders of magnitude faster. As shown in Figure 6, running more than 14 simulations on the simulator takes longer than training our meta-model when using the largest *radius*. On the other hand, when using the smallest *radius*, the time required to train the ANN is comparable to the time required to run around 70 simulations. Overall, this represents a significant improvement in terms of computation time and resources, and proves that the proposed approach supports not only accurate prediction, but computational time reduction as well, supporting efficient DSE of complex biological processes. All the experiments were performed on a 12th Gen Intel(R) Core(TM) i9-12900K with 24 threads and a maximum frequency of 5.2 GHz.

**Fig 6:**
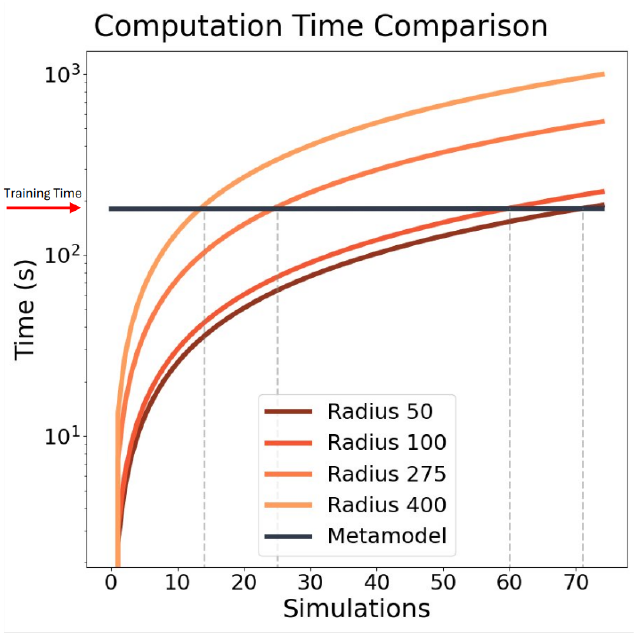
Computation time comparison between simulations with PhysiBoSS 2.0 and with our meta-model for all tested values of *tumor radius* (CPU time, in seconds). The initial offset of the meta-model time, as indicated by the red arrow, represents the training time (GPU time, in seconds). The CPU used for the simulations is a 12th Gen Intel(R) Core(TM) i9-12900K with 24 threads and a maximum frequency of 5.2 GHz, while the GPU used for the training of the model is an NVIDIA RTX A4000 with 16 GB of memory.

## V. Conclusions

In this paper, we proposed an LSTM-based surrogate modeling approach for multi-level simulations of complex biological processes applied to the simulation of TNF administration to a 3T3 mouse fibroblasts tumor spheroid based on PhysiBoSS 2.0. The meta-model accurately predicts the behavior of *N*_alive_, *N*_apoptotic_ and *N*_necrotic_, significantly reducing the simulation runtime by at least five orders of magnitude compared to the original simulator. The proposed meta-modeling approach is a valuable tool for computational biologists to harness the predictive power of multi-level hybrid model-based OvS while reducing the computational resources needed to perform a broad DSE. In this work, trainings based over four different experimental conditions, corresponding to four value of the *tumor radius*, generated four meta-models. In the future, we aim to build a more general meta-model encompassing these different scenarios. This will require to manage the different scales in the number of cells, dynamically adapting data normalization to the *tumor radius*.

Additionally, all simulations used for this use case and, consequently, all models’ predictions, are two-dimensional.

This reduces the time required for the dataset extraction but decreases the similarity to real in-vitro processes. To increase the expressivity and biological accuracy of the model, we aim to leverage PhysiBoSS 2.0 3D simulation to train the metamodel. In conclusion, this acceleration in DSE enables largescale, high-throughput explorations, making meta-modeling a powerful tool for identifying and optimizing effective treatment strategies and facilitating personalized medicine.

## Data availability

All the code employed in this study is publicly available on the GitHub repository at https://github.com/smilies-polito/LSTMeta-TNF.

## Acknowledgment

Marco Pietro Abrate acknowledges the London Interdisciplinary Doctoral Program (LIDO) and the Biotechnology and Biological Sciences Research Council (BBSRC) for funding his PhD scholarship. We would also like to thank Miguel Ponce de Leon, Vincent Noël, and Arnau Montagud for their insightful suggestions and feedback.

## References

[1] R. Bardini, G. Politano, A. Benso, and S. Di Carlo, “Multi-level and hybrid modelling approaches for systems biology,” Computational and structural biotechnology journal, vol. 15, 2017.

[2] Z. Ji, K. Yan, W. Li, H. Hu, and X. Zhu, “Mathematical and computational modeling in complex biological systems,” BioMed research international, vol. 2017, no. 1, 2017.

[3] J. M. Cardoso, J. G. F. Coutinho, and P. C. Diniz, “Chapter 8 - additional topics,” in Embedded Computing for High Performance, J. M. Cardoso, J. G. F. Coutinho, and P. C. Diniz, Eds. Boston: Morgan Kaufmann, 2017. [Online]. Available: https://www.sciencedirect.com/science/article/pii/B9780128041895000089

[4] J. V. S. do Amaral, J. A. B. Montevechi, R. de Carvalho Miranda, and W. T. de Sousa Junior, “Metamodel-based simulation optimization: A systematic literature review,” Simulation Modelling Practice and Theory, vol. 114, 2022.

[5] R. Bardini and S. Di Carlo, “Computational methods for biofabrication in tissue engineering and regenerative medicine-a literature review,” Computational and Structural Biotechnology Journal, 2024.

[6] A. Castrignanò, R. Bardini, A. Savino, and S. Di Carlo, “A methodology combining reinforcement learning and simulation to optimize the in silico culture of epithelial sheets,” Journal of Computational Science, vol. 76, 2024.

[7] L. Giannantoni, R. Bardini, and S. Di Carlo, “A methodology for cosimulation-based optimization of biofabrication protocols,” in International Work-Conference on Bioinformatics and Biomedical Engineering. Springer, 2022, pp. 179–192.

[8] M. Ponce-de Leon, A. Montagud, C. Akasiadis, J. Schreiber, T. Ntiniakou, and A. Valencia, “Optimizing dosage-specific treatments in a multi-scale model of a tumor growth,” Frontiers in Molecular Biosciences, vol. 9, 2022.

[9] J. Xu, E. Huang, L. Hsieh, L. H. Lee, Q.-S. Jia, and C.-H. Chen, “Simulation optimization in the era of industrial 4.0 and the industrial internet,” Journal of Simulation, vol. 10, 2016.

[10] R. Alizadeh, J. K. Allen, and F. Mistree, “Managing computational complexity using surrogate models: a critical review,” Research in Engineering Design, vol. 31, no. 3, 2020.

[11] E. D. Sozzo, D. Conficconi, A. Zeni, M. Salaris, D. Sciuto, and M. D. Santambrogio, “Pushing the level of abstraction of digital system design: A survey on how to program fpgas,” ACM Computing Surveys, vol. 55, no. 5, 2022.

[12] K. McBride and K. Sundmacher, “Overview of surrogate modeling in chemical process engineering,” Chemie Ingenieur Technik, vol. 91, no. 3, 2019.

[13] W. Fang, Y. Chen, and Q. Xue, “Survey on research of rnn-based spatiotemporal sequence prediction algorithms,” Journal on Big Data, vol. 3, no. 3, 2021.

[14] G. Letort, A. Montagud, G. Stoll, R. Heiland, E. Barillot, P. Macklin, A. Zinovyev, and L. Calzone, “Physiboss: a multi-scale agent-based modelling framework integrating physical dimension and cell signalling,” Bioinformatics, vol. 35, no. 7, 2019.

[15] L. Calzone, L. Tournier, S. Fourquet, D. Thieffry, B. Zhivotovsky, E. Barillot, and A. Zinovyev, “Mathematical modelling of cell-fate decision in response to death receptor engagement,” PLoS computational biology, vol. 6, no. 3, 2010.

[16] M. Ponce-de Leon, A. Montagud, V. Noïl, A. Meert, G. Pradas, E. Barillot, L. Calzone, and A. Valencia, “Physiboss 2.0: a sustainable integration of stochastic boolean and agent-based modelling frameworks,” npj Systems Biology and Applications, vol. 9, no. 1, 2023.

[17] A. Ghaffarizadeh, R. Heiland, S. H. Friedman, S. M. Mumenthaler, and P. Macklin, “Physicell: An open source physics-based cell simulator for 3-d multicellular systems,” PLoS computational biology, vol. 14, no. 2, 2018.

[18] G. Stoll, B. Caron, E. Viara, A. Dugourd, A. Zinovyev, A. Naldi, G. Kroemer, E. Barillot, and L. Calzone, “Maboss 2.0: an environment for stochastic boolean modeling,” Bioinformatics, vol. 33, no. 14, 2017.

[19] I. M. Gherman, Z. S. Abdallah, W. Pang, T. E. Gorochowski, C. S. Grierson, and L. Marucci, “Bridging the gap between mechanistic biological models and machine learning surrogates,” PLoS Computational Biology, vol. 19, no. 4, 2023.

[20] L. H. Gilpin, D. Bau, B. Z. Yuan, A. Bajwa, M. Specter, and L. Kagal, “Explaining explanations: An overview of interpretability of machine learning,” in 2018 IEEE 5th International Conference on data science and advanced analytics (DSAA). IEEE, 2018, pp. 80–89.

[21] M. Hwang, C. H. Leem, and E. B. Shim, “Toward a grey box approach for cardiovascular physiome,” The Korean Journal of Physiology & Pharmacology: Official Journal of the Korean Physiological Society and the Korean Society of Pharmacology, vol. 23, no. 5, 2019.

[22] Y. Zhao, C. Jiang, M. A. Vega, M. D. Todd, and Z. Hu, “Surrogate modeling of nonlinear dynamic systems: a comparative study,” Journal of Computing and Information Science in Engineering, vol. 23, no. 1, 2023.

[23] A. Sherstinsky, “Fundamentals of recurrent neural network (rnn) and long short-term memory (lstm) network,” Physica D: Nonlinear Phenomena, vol. 404, 2020.

[24] X. Song, Y. Liu, L. Xue, J. Wang, J. Zhang, J. Wang, L. Jiang, and Z. Cheng, “Time-series well performance prediction based on long short-term memory (lstm) neural network model,” Journal of Petroleum Science and Engineering, vol. 186, 2020.

[25] S. Hochreiter and J. Schmidhuber, “Long short-term memory,” Neural computation, vol. 9, no. 8, 1997.

[26] H. Kitano, “Systems biology: a brief overview,” science, vol. 295, no. 5560, 2002.

[27] S. Wang, K. Fan, N. Luo, Y. Cao, F. Wu, C. Zhang, K. A. Heller, and L. You, “Massive computational acceleration by using neural networks to emulate mechanism-based biological models,” Nature communications, vol. 10, no. 1, 2019.

[28] L. Burzawa, L. Li, X. Wang, A. Buganza-Tepole, and D. M. Umulis, “Acceleration of pde-based biological simulation through the development of neural network metamodels,” Current pathobiology reports, vol. 8, 2020.

[29] D. Bernard, A. Kobanda, and S. Cussat-Blanc, “Simulating cytotoxic t-lymphocyte and cancer cells interactions: An lstm-based approach to surrogate an agent-based model,” in International Symposium on Mathematical and Computational Oncology. Springer, 2021.

[30] P. Davidson, H. Trinh, S. Vekki, and P. Müller, “Surrogate modelling for oxygen uptake prediction using lstm neural network,” Sensors, vol. 23, no. 4, 2023.

[31] S. Li, G. Xie, J. Ren, L. Guo, Y. Yang, and X. Xu, “Urban pm2. 5 concentration prediction via attention-based cnn–lstm,” Applied Sciences, vol. 10, no. 6, 2020.

[32] Y. Tian, K. Zhang, J. Li, X. Lin, and B. Yang, “Lstm-based traffic flow prediction with missing data,” Neurocomputing, vol. 318, 2018.

[33] H. Bhatia, F. Aydin, T. S. Carpenter, F. C. Lightstone, P.-T. Bremer, H. I. Ingólfsson, D. V. Nissley, and F. H. Streitz, “The confluence of machine learning and multiscale simulations,” Current Opinion in Structural Biology, vol. 80, 2023.

[34] A. Paszke, S. Gross, F. Massa, A. Lerer, J. Bradbury, G. Chanan, T. Killeen, Z. Lin, N. Gimelshein, L. Antiga et al., “Pytorch: An imperative style, high-performance deep learning library,” Advances in neural information processing systems, vol. 32, 2019.

